# Somatic cell-derived BMPs induce male germ cell meiosis initiation during embryonic stage via regulating *Dazl* expression

**DOI:** 10.1101/2020.03.01.971937

**Authors:** Lianjun Zhang, Yaqiong Li, Yuqiong Hu, Limei Lin, Jingjing Zhou, Min Chen, Yan Qin, Yang Zhou, Min Chen, Xiuhong Cui, Fuchou Tang, Fei Gao

## Abstract

Germ cell fate is believed to be determined by the signaling from sexually differentiated somatic cell. However, the molecular mechanism remains elusive. In this study, ectopic initiation of meiosis in male germ cells was observed during embryonic stage by over-activating CTNNB1 in Sertoli cells. Somatic cell transcriptome and single germ cell RNA-seq analysis indicated that TGF-β signaling was activated after CTNNB1 over-activation. *In vitro* and *in vivo* experiments confirmed somatic cell-derived BMPs played crucial roles in germ cell meiosis initiation. Further studies revealed that *Dazl* was significantly increased in germ cells of CTNNB1 over-activated testes and induced by BMP signaling. DNMT3a and DNA methylation was also reduced in germ cells of CTNNB1 over-activated testes and increased by BMP signaling inhibitor treatment. Taken together, this study demonstrates that germ cell fate could be reprogrammed after sex determination. BMP signaling pathway is involved in germ cell meiosis initiation via up-regulating *Dazl* expression.

## Introduction

Generation of haploid gametes via meiosis is a unique property of germ cells and essential for sexual reproduction (McLaren, 1984, 2001). Whether primordial germ cells (PGCs) develop as oogonia or pro-spermatogonia depending on the differentiation of somatic cells during sex determination (Adams and McLaren, 2002; McLaren, 1991). Briefly, *Sry* is transiently expressed in male somatic cells between E10.5 and E12.5, which directs Sertoli cells differentiation (Albrecht and Eicher, 2001). Germ cells in male gonads are arrested in G0/G1 stage after several divisions during embryonic stage and re-enter cell cycle to initiate meiosis after birth (Western et al., 2008). On the other hand, R-spondin1 (RSPO1)/CTNNB1 pathway promotes the differentiation of granulosa cells in female gonads (Chassot et al., 2008; Tomizuka et al., 2008) and female germ cells enter meiosis directly following granulosa cells differentiation, and then arrested at the diplotene stage of prophase I (Borum, 1961).

Up to now, RA (retinoic acid) is considered as the most important extrinsic factor which stimulates germ cells to enter meiosis by inducing the expression of STRA8 (Stimulated by Retinoic Acid 8) and REC8 (REC8 meiotic recombination protein) in germ cells (Bowles et al., 2006; Koubova et al., 2014; Koubova et al., 2006). STRA8 is required for pre-meiotic DNA replication (Baltus et al., 2006) and REC8 is essential for sister chromatid’s separation (Xu et al., 2005). As an intrinsic factor, DAZL (deleted in azoospermia-like) is strictly expressed in germ cells and required for the “license” of germ cell meiosis initiation (Lin et al., 2008; Lin and Page, 2005). The expression of *Stra8* and *Rec8* are markedly reduced in germ cells of *Dazl*-deficient mice, suggesting that *Dazl*-deficient germ cells don’t respond to RA induction and initiate meiosis (Lin and Page, 2005).

It’s believed that germ cells enter meiosis spontaneously during embryonic stage unless are specifically prevented by meiosis-inhibiting factors. CYP26B1 (Cyp26 family of cytochrome P450 oxidase 1) is first identified as a meiosis inhibitor which is abundantly expressed in Sertoli cells during embryonic stage and catalyzes the oxidization of RA to inactive forms (MacLean et al., 2001). Inactivation of CYP26B1 leads to the ectopic initiation of meiosis in male germ cells during embryonic stage (Bowles et al., 2006; MacLean et al., 2007). Another meiosis-inhibiting substance is FGF9 (fibroblast growth factor 9) also produced by Sertoli cells, which acts antagonistically with RA to determine male germ cell fate (Bowles et al., 2010). Male-to-female sex reversal is observed in *Fgf9*-null mice, and could be rescued by *Wnt4* (*wingless-type MMTV integration site family, member 4*) deletion (Jameson et al., 2012a). FGF9 also acts directly on germ cells by upregulating RNA binding protein NANOS2 (nanos C2HC-type zinc finger 2) (Bowles et al., 2010). *Nanos2* is male meiotic gatekeeper specifically expressed in germ cells and plays pivotal roles in germ cell sexual differentiation (Saga, 2010; Suzuki and Saga, 2008). Inactivation of *Nanos2* results in meiotic initiation in male germ cells and upregulation of oocyte differentiation associated genes (Suzuki et al., 2010; Suzuki et al., 2012). Inversely, female germ cells fail to enter meiosis and start expressing male-specific genes after NANOS2 over-expression (Suzuki and Saga, 2008).

Several studies have demonstrated that TGF-β signaling is also involved in regulating meiosis and germ cell development (Spiller et al., 2017; Wu et al., 2013). Nodal/activin pathway is activated in both male germ cells and somatic cells which induces *Nanos2* expression. Disruption of Nodal/Activin signaling leads to male germ cells meiosis and increased expression of female-specific genes in somatic cells (Souquet et al., 2012; Spiller et al., 2013; Tassinari et al., 2015). Moreover, deletion of *Smad4* in germ cells results in female germ cells meiosis defect (Wu et al., 2016). Moreover, recent study demonstrates that BMPs (bone morphogenetic proteins) direct the female fate determination of PGCs/PGC-like cells *in vitro* induction (Miyauchi et al., 2017).

Our previous study demonstrates that over-activation of CTNNB1 in Sertoli cells results in Sertoli to granulosa-like cells transformation (Li et al., 2017). Interestingly, in this study, we find the ectopic initiation of meiosis in male germ cells of CTNNB1 overactivated mice during embryonic stage. Further studies demonstrate that somatic cell derived BMPs plays essential roles in germ cell meiosis initiation. Moreover, BMP signaling pathway involves in orchestrating the germ cell meiosis most likely via up-regulating *Dazl* expression and it is probably mediated by repressing DNA methylation.

## Results

### Ectopic expression of meiotic genes in germ cells of Ctnnb1_+/flox(ex3)_ AMH-Cre mice during embryonic stage

Our previous studies have demonstrated that over-activation of *Ctnnb1* by deletion of exon 3 results in testicular cord disruption and Sertoli to granulosa-like cells transformation (Chang et al., 2008; Li et al., 2017). To track the differentiation of germ cells in *Ctnnb1_+/flox(ex3)_ AMH-Cre* mice (hereafter referred as COA mice unless otherwise specified), the expression of germ cell specific and meiosis-associated genes was analyzed by immunofluorescence and quantitative RT-PCR. Interestingly, several meiotic markers were expression in the germ cells of COA testes. No STRA8 was observed in germ cells of control testes from E13.5 to E17.5 (Fig. S1Aa-e, white arrows). By contrast, it was detected in a large number of germ cells in COA testes from E13.5 to E16.5 (Fig. S1Af-i, white arrowheads), which is a period associated character in germ cells of control ovaries during embryonic stage (Fig. S1Ak-m, white arrows). γH2AX, a marker of DNA double-strand breaks (DSBs), was also detected in germ cells of COA testes (Fig. S1Bg-j, white arrowheads) from E14.5 to E17.5 which was similar to the germ cells in control ovaries (Fig. S1Bk-o, white arrows). As expected, no γH2AX was detected in control testes (Fig. S1Ba-e, white arrows). Hematoxylin and eosin (H&E) staining also showed that the chromatin was highly compacted in germ cells of COA mice (Fig. S2D-F, white arrowheads) but not in control male germ cells (Fig. S2A-C, white arrows), and the similar chromatin morphology was observed in control female germ cells (Fig. S2G-I, white arrowheads).

Importantly, synaptonemal complex protein SYCP3, a typical marker of meiosis was also abundantly expressed in the germ cells of COA testes. As shown in Figure 1 A, thread-like SYCP3 signal was detected in germ cells of COA testes at E16.5 and E17.5 (Fig. 1Ai, j, white arrowheads), which was restrictedly expressed in the germ cells of control ovaries during embryonic stage (Fig. 1Ak-o, white arrowheads), but not in the germ cells of control testes (Fig. 1Aa-e, white arrows). To further determine the meiotic stages of germ cells in COA testes, chromosome spreads of meiocytes were prepared and stained with SYCP3 antibody. As shown in Figure 1B, germ cells at leptotene, zygotene, pachytene, and diplotene stage of meiosis I were observed in COA testes which was normally detected in germ cells of control ovaries during embryonic stage (Fig. 1Ba). The numbers of germ cells at pachytene and diplotene stages in COA testes was less than that in control ovaries (Fig. 1Bb, c). The mRNA level of meiotic genes including *Stra8, Dmc1*, *Rec8, Sycp3* and *Sycp1* at E16.5 was also analyzed by quantitative RT-PCR and was significantly increased in COA testes compared to that of control testes, including the intrinsic factor *Dazl* (Fig. 1Cb). Male germ cell specific gene (*Dnmt3l*) and pluripotency genes (*Oct4* and *Sox2*) were decreased in COA testes, whereas female germ cell specific gene *Foxo3* and fetal oogenesis-specific gene *Sohlh2* was increased in COA testes (Fig. 1Ca). Collectively, all these results indicated that germ cells in COA testes ectopically initiated meiosis during embryonic stage.

**Figure 1.**
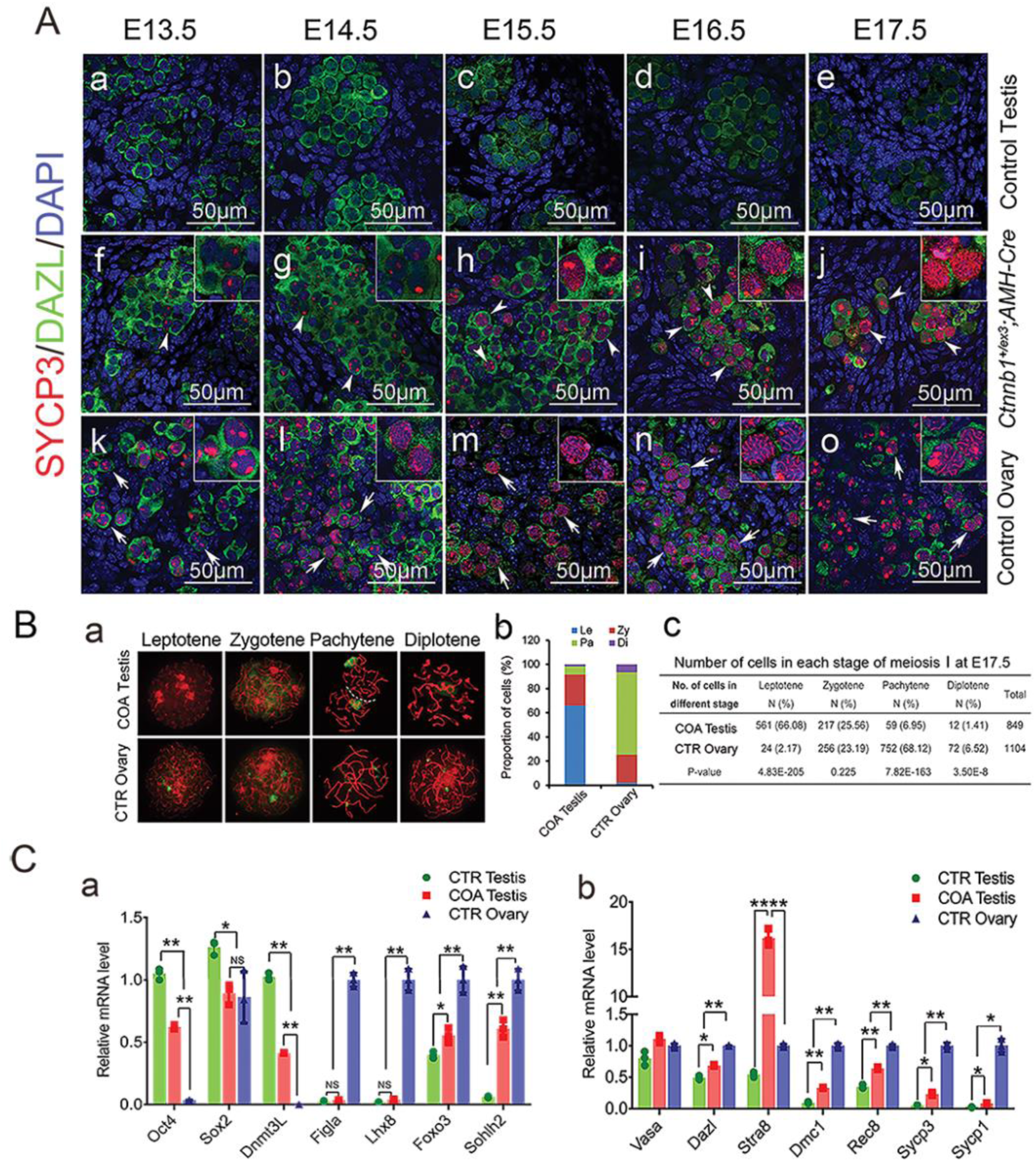
Abnormal initiation of meiosis in germ cells of *Ctnnb1_+/flox(ex3)_ AMH-Cre* mice during embryonic stage. **A**. SYCP3 protein was detected in the germ cells of *Ctnnb1_+/flox(ex3)_ AMH-Cre* testes during embryonic stage. No SYCP3 signal was detected in DAZL-positive germ cells (green, white arrows) of control testes from E13.5 to E17.5 (Aa-Ae). In *Ctnnb1_+/flox(ex3)_ AMH-Cre* testes, the foci of SYCP3 protein (red) was noted in a small portion of germ cells at E13.5 (Af, white arrowheads) and E14.5 (Ag, white arrowheads), and thread-like SYCP3 signal was detected at E15.5 and E17.5 (Ah-Aj, white arrowheads). In control ovaries, the foci of SYCP3 protein were noted in germ cells at E13.5 (Ak, white arrows). Thread-like SYCP3 signal was detected in germ cells from E14.5 to E17.5 (Al-Ao, white arrows). **B**. Chromosome spreads of germ cells in control ovaries and *Ctnnb1_+/flox(ex3)_ AMH-Cre* testes. Representative images of synaptonemal complex at Leptotene, Zygotene, Pachytene, and Diplotene stages of meiosis (Ba). Percentage (Bb) and number (Bc) of meiotic germ cells at different stages of meiosis prophase I. Le, Leptotene; Zy, Zygotene; Pa, Pachytene; Di, Diplotene. **C**. The mRNA level at E16.5 gonads was examined by quantitative RT-PCR. Pluripotency genes *Oct4*, *Sox2* and male germ cell marker *Dnmt3L* was decreased in *Ctnnb1_+/flox(ex3)_ AMH-Cre* testes; whereas the expression of embryonic female markers *Foxo3* and *Sohlh2* was increased in *Ctnnb1_+/flox(ex3)_ AMH-Cre* testes compared to control testes (Ca). Meiotic genes (*Dazl*, *Stra8*, *Dmc1*, *Rec8*, *Scp3*, *Scp1*) were significantly increased in *Ctnnb1_+/flox(ex3)_ AMH-Cre* testes compared to control testes (Cb). CTR, control; COA, *Ctnnb1_+/flox(ex3)_ AMH-Cre*. Data are presented as the mean ± SD. Two-way ANOVA analysis was used to test significance (*P< 0.05; **P< 0.01). See also Figure S1, S2.

### Somatic cell-derived secreted factors played essential roles in inducing germ cell meiosis initiation

It has been reported that male germ cells are prevented to enter meiosis by Sertoli cell-expressed CYP26B1 during embryonic stage (Bowles et al., 2006; Koubova et al., 2006; MacLean et al., 2001). To test whether the ectopic meiosis initiation in germ cells of COA testes is due to the testicular cord disruption, the expression of meiotic genes in germ cells of *DTA_+/flox_ AMH-Cre* and *Wt1_+/flox_ AMH-Cre* mice were examined by immunostaining. In *DTA_+/flox_ AMH-Cre* mice, Sertoli cells were ablated by Diphtheria toxin upon Cre activation (Brockschnieder et al., 2004; Rebourcet et al., 2018). The testicular cords were disrupted in *Wt1_+/flox_ AMH-Cre* mice due to the Sertoli to Leydig-like cell transformation (Chang et al., 2008; Zhang et al., 2015a). However, no STRA8 and γH2AX were detected in the germ cells of these two mouse models at E16.5 (Fig. S3D, E, G, H), only very weak SYCP3 was observed (Fig. S3F, I). These results indicated that the disruption of testicular cords does not cause ectopic meiosis initiation in male germ cells during embryonic stage.

Given the fact that Sertoli cells were transformed into granulosa-like cells in COA testes, the abnormal meiosis initiation of germ cells is probably induced by the factors derived from granulosa-like cells. To test this hypothesis, germ cells (GCs) and somatic cells (SCs) from E13.5 control ovaries and testes were re-aggregated and cultured *in vitro*. The differentiation of germ cells was examined after 3 days culture (Fig. S4A). As shown in Figure S4B, germ cells with thread-like SYCP3 were observed in (GCs♀+SCs♀) aggregated tissue (Fig. S4Ba). Interestingly, thread-like SYCP3 was also noted in male germ cells (GFP+) aggregated with female somatic cells (Fig. S4Bb). By contrast, only very weak or dot-like SYCP3 was detected in male germ cells aggregated with male somatic cells (Fig. S4Bc). These results suggested that the factors derived from female somatic cells were essential for germ cell meiosis and the ectopic meiosis initiation in the gem cells of COA testes was most likely triggered by the factors derived from the transformed granulosa-like cells.

### TGF-β signaling pathway was activated both in Sertoli cells and germ cells of Ctnnb1_+/flox(ex3)_ AMH-Cre testes

To explore the underlying mechanism which leads to the ectopic initiation of meiosis in COA germ cells, RNA-sequencing assay was performed with isolated Sertoli cells from control and COA mice at E13.5 and E14.5. We first profiled the upregulated and downregulated genes in COA Sertoli cells compared to control Sertoli cells at E13.5 and E14.5 (Fig. 2A). Of note, 846 genes were significantly changed in COA Sertoli cells with 374 genes were up-regulated and 472 genes were down-regulated (log2 FC > 1) compared to control Sertoli cells (Fig. 2B and C). Results of KEGG analysis revealed that the most differentially expressed genes were enriched in TGF-β and WNT signaling pathway (Fig. 2D). Further analysis showed that the ligands in BMP signaling pathway (e.g. *Bmp2*, *Bmp4*, *Bmp5*) were significantly upregulated in Sertoli cells of COA mice (Fig. 2E). The ligands in WNT signaling pathway (e.g. *Wnt10b*, *Wnt11*, *Wnt2*, *Wnt4*, *Wnt5b*, *Wnt7b*, *Wnt9a*) was also upregulated in Sertoli cells of COA mice (Fig. 2F).

**Figure 2.**
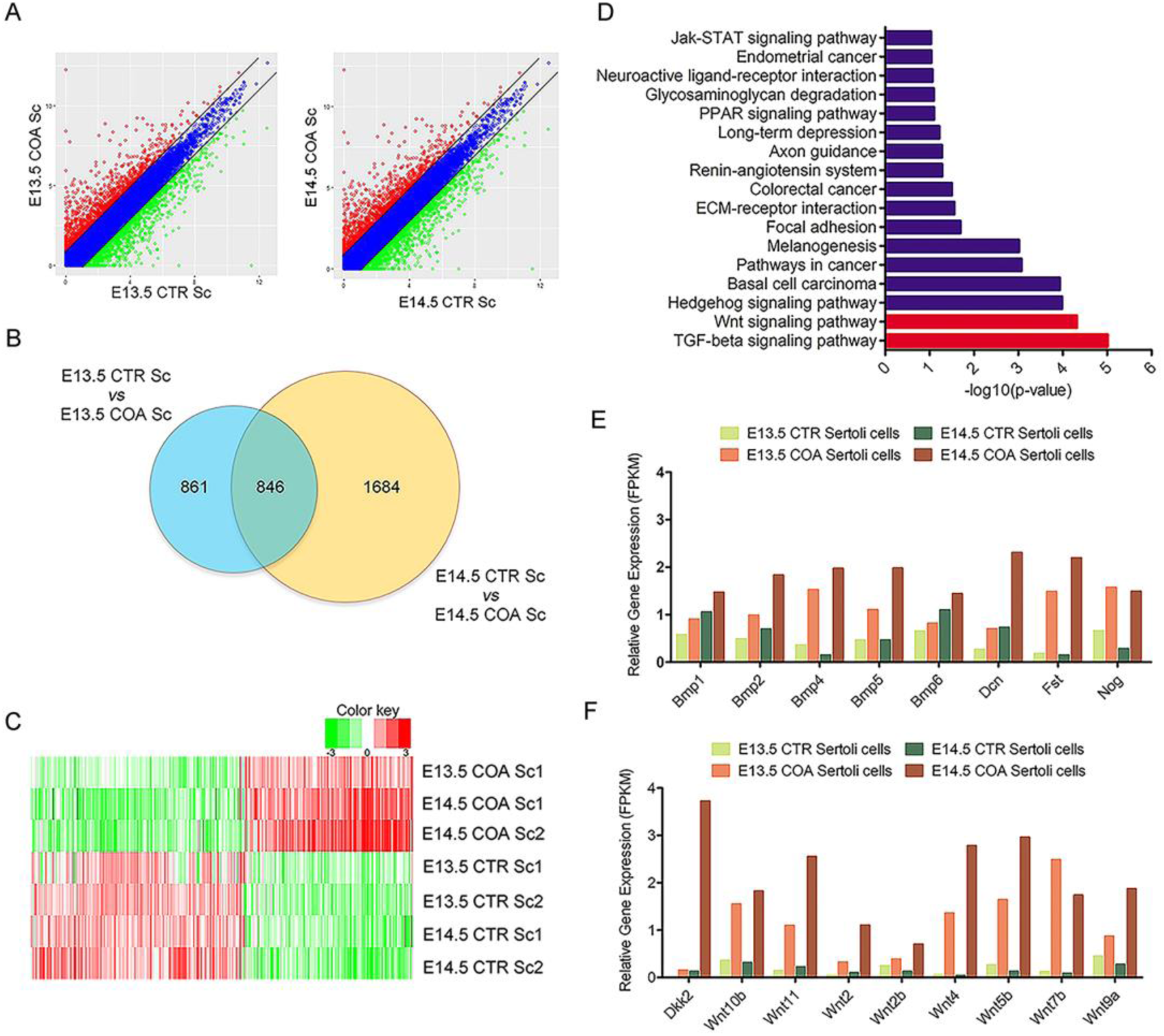
Differentially expressed genes in Sertoli cells of *Ctnnb1_+/flox(ex3)_ AMH-Cre* testes. **A**. Scatter-plot analysis of differentially expressed genes (DEGs) between COA Sertoli cells (Sc) and CTR Sertoli cells at E13.5 (left) and E14.5 (right). Red dots represented up-regulated genes and green dots represented downregulated genes in COA Sertoli cells versus CTR Sertoli cells. **B**. Venn diagram of numbers of DEGs between COA and CTR Sertoli cells at E13.5 and E14.5. Total 846 genes (log2 FC > 1) were differentially expressed at both E13.5 and E14.5. **C**. Heat maps of all the 846 DEGs in control and COA Sertoli cells. **D**. KEGG analysis of the DEGs. **E**. Relative gene expression level of DEGs in TGF-β signaling pathway. **F**. Relative gene expression level of DEGs in WNT signaling pathway. CTR, control; COA, *Ctnnb1_+/flox(ex3)_ AMH-Cre*.

To identify the differentially expressed genes in germ cells responding to the signals from somatic cells, single-cell RNA sequencing assay was performed with isolated germ cells from control female, control male and COA male mice at E14.5 and E15.5. Results of principal component analyses (PCA) showed that the gene profile of germ cells from COA mice was separated from control male and control female germ cells both at E14.5 and E15.5 (Fig. 3A). The results of clustered heatmap analysis of 800 differentially expressed genes based on PCA was shown in Figure 3B. Then we further filtrated 467 genes which were differentially expressed at least 2-fold change in both COA and control female germ cells compared to control male germ cells at E14.5 and E15.5 (Fig. 3C). KEGG pathway analysis of these genes showed that metabolic pathways were most significantly changed. As expected, TGF-β signaling pathway was also enriched along with cell cycle and oocyte meiosis pathway (Fig. 3D). Relative expression of meiotic genes (such as *Dazl*, *Stra8*, *Spo11*, *Rec8*, *Dmc1* and *Sycp3*) was shown in Figure 3E to confirm the meiotic state of isolated germ cells. Both *Stra8* and *Rec8* were dramatically upregulated in COA germ cells compared to control male and female germ cells at E15.5 and *Dmc1* and *Sycp3* was increased in COA germ cells compared to male germ cells at E15.5. Interestingly, *Dazl* expression was up-regulated in germ cells of COA mice compared to control male germ cells which was comparable to that in control female germ cells at E15.5 (Fig. 3E). The represented up-regulated downstream genes of TGF-β signaling pathway (*Bmpr1a*, *Ep300*, *E2f4*, *E2f5*, *Smurf2*, *Rbx1*, and *Tfdp1*) in COA mice were shown in Figure 3F.

**Figure 3.**
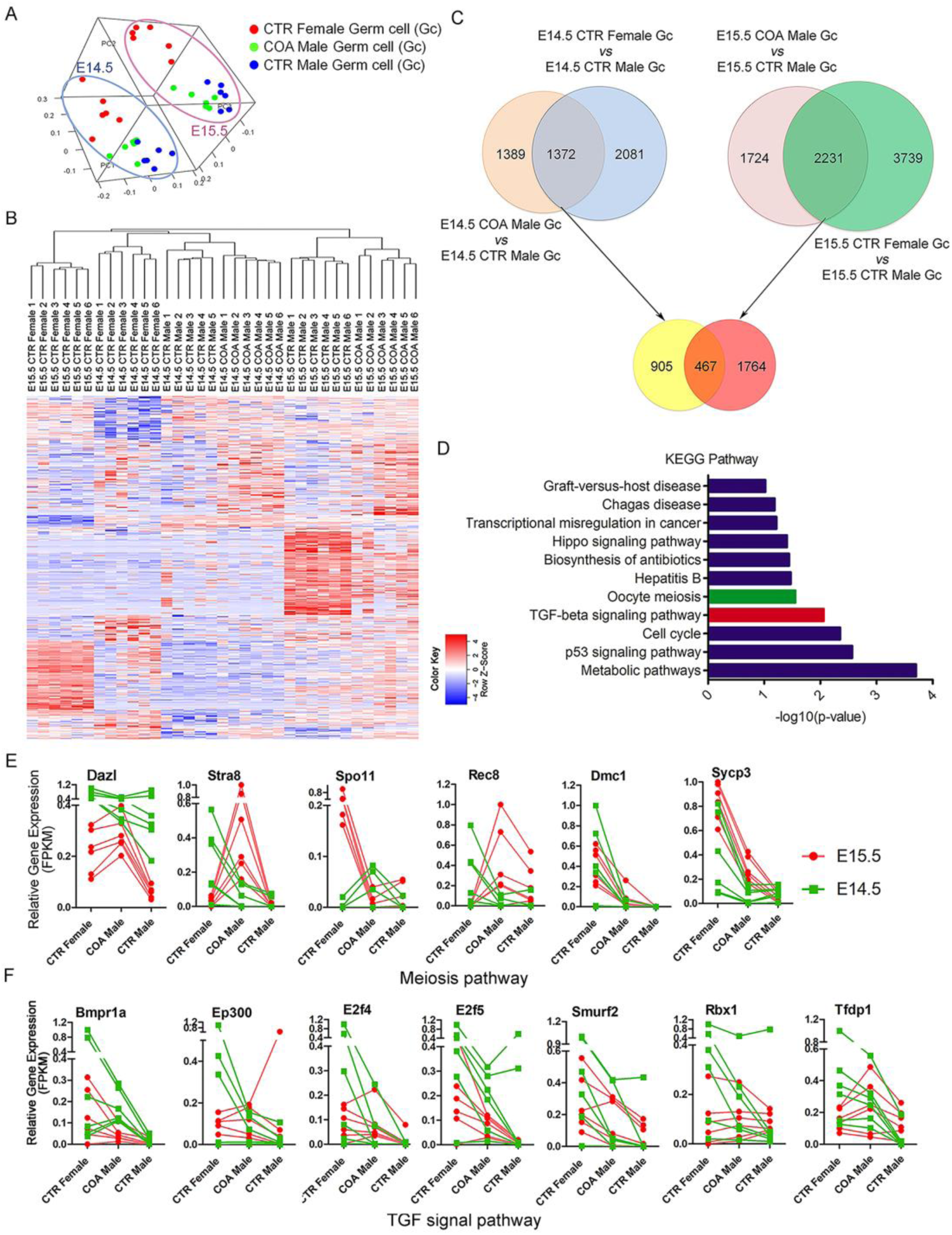
Single-cell RNA sequencing analysis of germ cells from control female, control male and COA male mice at E14.5 and E15.5. **A**. Principal component analyses (PCA) of germ cells from control female, control male and COA male mice. **B**. Heat maps of 800 DEGs based on PCA analysis in germ cells of control male, COA male and control female mice at E14.5 and E15.5. **C**. Venn diagram of numbers of DEGs in germ cells from control male, COA male and control female mice at E14.5 and E15.5. **D**. KEGG pathway analysis of 467 DEGs. **E**. The differential expression of meiotic marker genes. **F**. The differential expression of genes in TGF-β signaling pathway. CTR, control; COA, *Ctnnb1_+/flox(ex3)_ AMH-Cre*.

### BMP signaling pathway and RA synergistically induced germ cell meiosis initiation

To further test whether BMP and WNT signaling pathways are involved in meiosis initiation of COA mice, testes from E13.5 embryos were cultured *in vitro* and treated with RA, BMP signaling inhibitor LDN-193189, and WNT signaling inhibitor XAV-939. After 3 days culture, SYCP3 was examined by immunostaining. As shown in Fig. 4A, scattered SYCP3 foci were observed in the germ cells of control testes with RA treatment (Fig. 4Aa, g). Thread-like SYCP3 signal were detected in the germ cells of COA testes with RA treatment (Fig. 4Ac, i), whereas only scattered SYCP3 foci were noted in the germ cells without RA treatment (Fig. 4Ab, h). Interestingly, the expression of SYCP3 in the germ cells of RA treated COA testes was significantly reduced with dot-like SYCP3 signal after LDN-193189 and XAV-939 treatment (Fig. 4Ad-f, j-l). The mRNA level of meiotic genes was also significantly decreased in COA testes after LDN-193189 and XAV-939 treatment (Fig. 4B). These results suggested that RA is essential for meiosis initiation but not sufficient. To further address whether BMP and WNT signaling are also involved in control female germ cells meiosis initiation, the genital ridges from E11.5 control female embryos were cultured *in vitro* and treated with XAV-939 and LDN-193189. As shown in Fig 4C, thread-like SYCP3 signal were detected in germ cells of control ovaries with DMSO treatment (Fig. 4Ca, d), whereas only scattered foci of SYCP3 signal were detected after LDN-193189 treatment (Fig. 4Cb, e), and the signal of SYCP3 was decreased with XAV treatment (Fig. 4Cc, f). Consistently, the mRNA level of other meiotic genes was also significantly decreased after LDN-193189 treatment (Fig 4D). These results suggested that both BMP and WNT signaling are involved in the meiosis initiation of germ cells in COA testes, and BMP signaling plays more important roles in this process.

**Figure 4.**
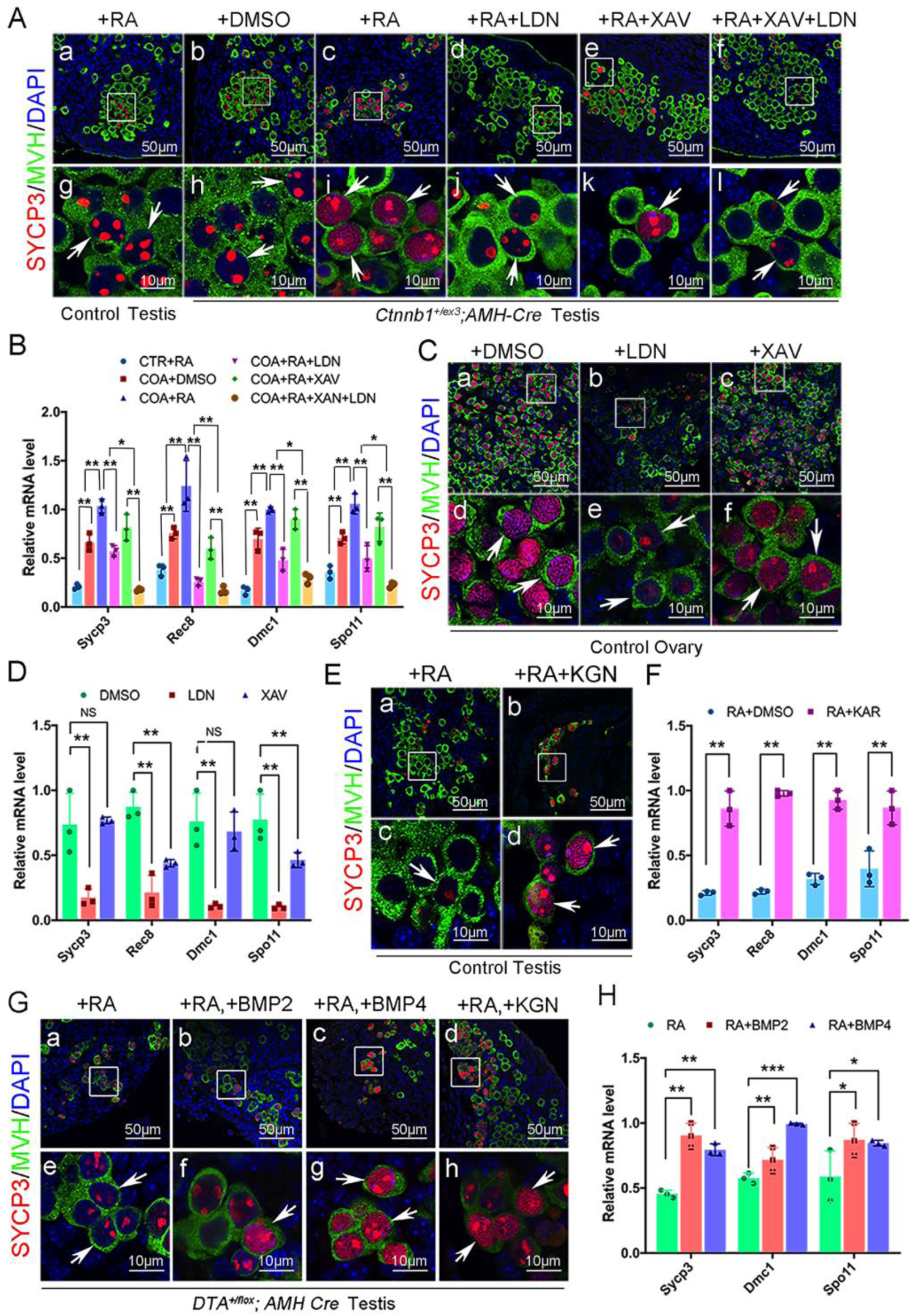
BMP signaling pathway was required for meiosis initiation of germ cells. The gonads were cultured *in vitro* for 3 days. **A.** Meiosis of germ cells in COA testes was inhibited by LDN-193189 and XAV-939 treatment. Scattered SYCP3 foci were observed in the germ cells of control testes with RA treatment (Aa, Ag) and in the germ cells of COA testes without RA treatment (Ab, Ah), whereas thread-like SYCP3 signal was detected in the germ cells of COA testes with RA treatment (Ac, Ai). Dot-like SYCP3 signal were observed in the germ cells of COA testes after LDN-193189 and XAV-939 treatment (Ad, Aj). **B.** The relative mRNA level of meiosis-related genes was examined by quantitative RT-PCR. **C.** The progress of meiosis in normal ovaries was repressed by LDN-193189 treatment. Thread-like SYCP3 signal were detected in germ cells of control ovaries with DMSO treatment (Ca, Cd), whereas only scattered foci of SYCP3 signal were detected after LDN-193189 treatment (Cb, Ce). The number of germ cells with thread-like SYCP3 signal was significantly decreased with XAV treatment (Cc, Cf). **D.** The relative mRNA level of meiosis-related genes in ovaries was examined by quantitative RT-PCR. **E.** Meiosis in CTR testes was promoted by treatment with BMP signaling activator Kartogenin. Weak SYCP3 signal was noted in germ cells of control testes with RA only treatment (Ea, Ec), and a small portion of germ cells with thread-like SYCP3 signal was noted in control testes with RA+Kartogenin (KGN) treatment (Eb, Ed). **F.** The mRNA level of meiosis-related genes was significantly increased in RA+Kartogenin treated testes. **G.** Meiosis of germ cells in *DTA_+/flox_ AMH-Cre* testes was induced by treatment of BMPs and activator Kartogenin. Dot-like SYCP3 signal was noted in the germ cells of *DTA_+/flox_ AMH-Cre* testes with RA only treatment (Ga, Ge). The germ cells with thread-like SYCP3 signal were observed in in the germ cells of *DTA_+/flox_ AMH-Cre* testes with RA+BMP2 (Gb, Gf), RA +BMP4 (Gc, Gg) and RA+Kartogenin (Gd, Gh) treatment. **H.** The mRNA level of meiosis-related genes was examined by quantitative RT-PCR. CTR, control; COA, *Ctnnb1_+/flox(ex3)_ AMH-Cre*. Data are presented as the mean ± SD. Two-way ANOVA and the student ‘s unpaired t-test analysis was used to test significance (*P< 0.05; **P< 0.01; ns: no significance).

The function of BMPs in germ cell meiosis was further examined using *in vitro* cultured E13.5 control and *DTA_+/flox_ AMH-Cre* testes and treated with RA, BMPs, and smad4/smad5 pathway activator Kartogenin (KGN). As shown in Figure 4E, germ cells with thread-like SYCP3 were detected in testes with RA+KGN treatment (Fig. 4Eb, d), whereas only scattered SYCP3 foci were observed in germ cells of RA only treated testes (Fig. 4Ea, c). Results of quantitative RT-PCR also showed that the mRNA level of meiotic genes was significantly increased in germ cells with RA+KGN treatment (Fig. 4F). For *DTA_+/flox_ AMH-Cre* testes, dot-like SYCP3 was noted in the germ cells with RA only treatment (Fig. 4Ga, e), whereas some germ cells with thread-like SYCP3 signal were observed after RA+BMP2 or RA+BMP4 treatment (Fig. 4Gb, f and c, g), and RA+Kartogenin (Fig. 4Gd, h) treatment. Moreover, the mRNA level of meiotic genes was also significantly increased after BMP2 and BMP4 treatment (Fig. 4H).

To further confirm the function of BMP signaling pathway in germ cell meiosis, pregnant female mice were intraperitoneally injected with LDN-193189 every 12 hours from E13.5 to E16.5 and the expression of SYCP3 was examined by immunostaining at E16.5 (Fig. 5A). As shown, no SYCP3 signal was detected in the germ cells of control testes with (Fig. 5Bk, l) or without (Fig. 5Bi, j) LDN-193189 injection. Thread-like SYCP3 signal were observed in the germ cells of control ovaries (Fig. 5Ba, b) and COA testes (Fig. 5Be, f) without LDN-193189 injection. However, only scattered foci of SYCP3 was detected in the germ cells of control ovaries (Fig. 5Bc, d) and COA testes (Fig. 5Bg, h) with LDN-193189 injection. These results suggested that BMP signaling pathway is essential for germ cell meiosis initiation and the ectopic initiation of meiosis in germ cells of COA testes is most likely induced by the BMPs derived from transformed Sertoli cells.

**Figure 5.**
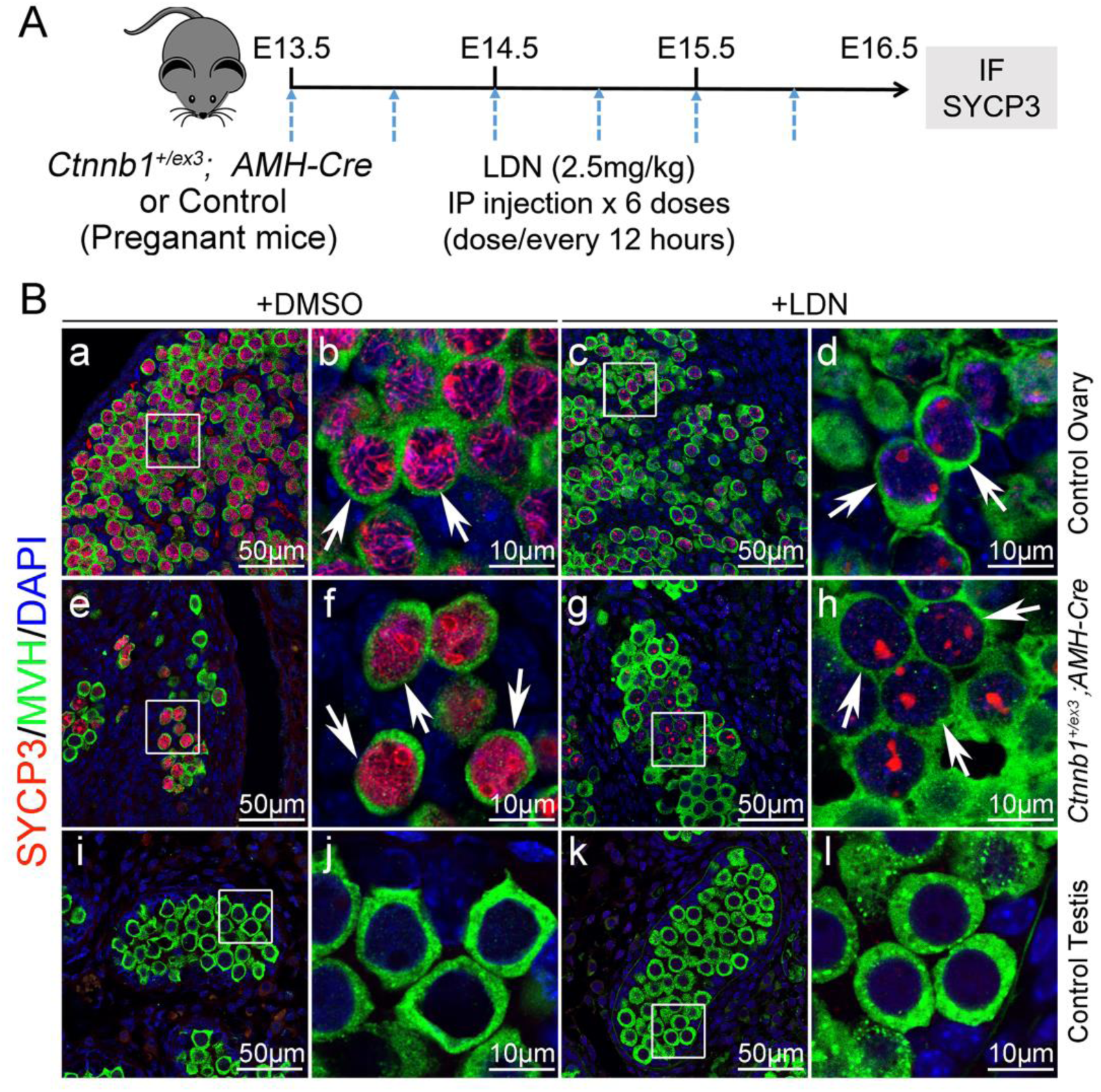
The meiosis of germ cells in *Ctnnb1_+/flox(ex3)_ AMH-Cre* testes and control ovaries was repressed by intraperitoneal injection of BMP signaling inhibitor LDN-193189. The expression of SYCP3 in germ cells of control ovaries, control testes, and *Ctnnb1_+/flox(ex3)_ AMH-Cre* testes at E16.5 was examined by immunostaining. **A.** The schematic diagram of intraperitoneal (IP) injection. **B.** LDN-193189 inhibited the progress of meiosis both in *Ctnnb1_+/flox(ex3)_ AMH-Cre* testes and CTR ovaries. Thread-like SYCP3 signal were observed in the germ cells of control ovaries (Ba, Bb) and *Ctnnb1_+/flox(ex3)_ AMH-Cre* testes (Be, Bf) with DMSO injection. Only scattered foci of SYCP3 signal were observed in the germ cells of control ovaries (Bc, Bd) and *Ctnnb1_+/flox(ex3)_ AMH-Cre* testes (Bg, Bh) with LDN-193189 treatment for 3 days. No SYCP3 signal was detected in the germ cells of control testes with (Bk, Bl) or without (Bi, Bj) LDN-193189 injection. CTR, control; COA, *Ctnnb1_+/flox(ex3)_ AMH-Cre*.

### The expression of Dazl was induced by BMP signaling pathway

*Dazl* is the intrinsic factor which is indispensable for RA response and meiosis initiation (Lin et al., 2008). In this study, we found that the expression of *Dazl* was significantly increased with RA+KAR treatment but not with RA treatment in E13.5 cultured testes (Fig. S5A). The expression of *Dazl* at E13.5 in *DTA_+/flox_ AMH-Cre* testes was also induced by BMP2 and BMP4 treatment (Fig. S5B). By contrast, the expression of *Dazl* in control ovaries was significantly decreased with LDN treatment, but not with XAV treatment (Fig. S5C). Moreover, the mRNA level of *Dazl* in RA treated COA testes was significantly higher than that in RA treated control testes, which was significantly decreased after LDN and XAV treatment (Fig. S5D). All together, these results indicated that the expression of *Dazl* was induced by BMP signaling pathway and BMPs involved in germ cell meiosis most likely via inducing *Dazl* expression.

### DNA methylation and DNMT3a was repressed by BMP signaling pathway in germ cells of COA testes

DNA methylation in germ cells is globally erased at ∼ E9.5 in both male and female embryos in mouse model (Hill et al., 2018). It is re-established in male germ cells at around E15.5, but much later in female germ cells (Smallwood and Kelsey, 2012). The results of gene ontology analysis of single germ cell RNA-sequencing data at E15.5 revealed that DNA methylation was enriched (Fig. S6A) and the expression of DNA methyltransferase *Dnmt3a* and *Dnmt3l* was decreased in germ cells of COA mice compared to control at E15.5 (Fig. S6Ba, c). Immunostaining results showed that 5mC was significantly decreased in germ cells of COA testes at E15.5 compared to control male germ cells, which was comparable to that of female germ cells (Fig. S6Ca-c). The expression of DNMT3a and DNMT3L was significantly decreased in germ cells of COA testes (Fig. S6Ce, n), whereas only DNMT3a was consistent with RNA-sequence result and comparable to control female germ cells.

To test whether DNA methylation is regulated by RA and BMP signaling, 5mC level and the expression of DNMT3a were examined by immunostaining in control and COA testes. As shown, both 5mC and DNMT3a were detected in germ cells of control testes (Fig. 6A and B a, g, m) and not repressed by RA treatment (Fig. 6A and Bb, h, n). However, 5mC and DNMT3a were not detected in germ cells of RA treated COA testes (Fig. 6A and B c, i, o), whereas it was significantly increased after LDN and XAV+LDN treatment (Fig. A and B d, j, p, f, l, r). Additionally, the level of 5mC and DNMT3a was not increased in COA germ cells with RA+XAV treatment (Fig. 6A and Be, k, q). These results indicated that DNA methylation in germ cells of COA testes is repressed by BMP signaling, and which is probably via down-regulating *Dnmt3a* expression.

**Figure 6.**
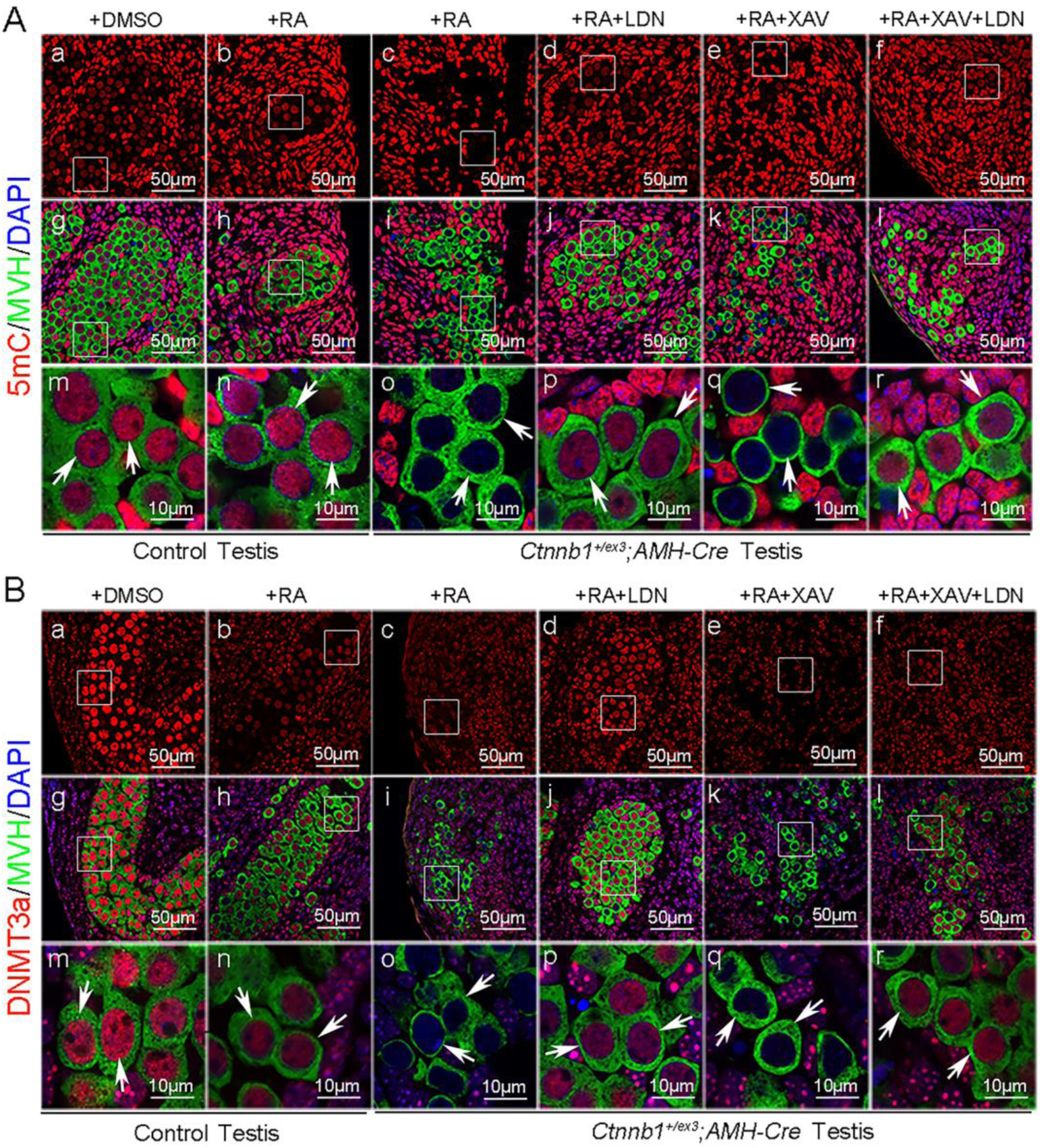
DNA methylation in germ cells of *Ctnnb1_+/flox(ex3)_ AMH-Cre* testes was repressed by BMP signaling pathway. Testes from control and *Ctnnb1_+/ flox (ex3)_ AMH-Cre* embryos at E13.5 were cultured *in vitro* and treated with RA, BMP signaling inhibitor LDN-193189, and WNT signaling inhibitor XAV-939. The level of 5mC and expression of DNMT3a were examined by immunostaining. **A.** High level of 5mC was detected in the germ cells of both DMSO (Aa, Ag, Am) and RA (Ab, Ah, An) treated control testis. No 5mC signal was detected in the germ cells of *Ctnnb1_+/ flox (ex3)_ AMH-Cre* testes treated with RA (Ac, Ai, Ao). The level of 5mC was significantly increased in the germ cells of *Ctnnb1_+/ flox (ex3)_ AMH-Cre* testes after LDN-193189 (Ad, Aj, Ap) and LDN+XAV (Af, Al, Ar) treatment, whereas it was not changed with XAV-939 treatment (Ae, Ak, Aq). **B.** High level of DNMT3a expression was detected in the germ cells of both DMSO (Ba, Bg, Bm) and RA (Bb, Bh, Bn) treated control testis. No DNMT3a signal was detected in the germ cells of *Ctnnb1_+/ flox (ex3)_ AMH-Cre* testes treated with RA (Bc, Bi, Bo). The expression of DNMT3a was significantly increased in the germ cells of *Ctnnb1_+/ flox (ex3)_ AMH-Cre* testes after LDN-193189 (Bd, Bj, Bp) and LDN+XAV (Bf, Bl, Br) treatment, whereas it was not changed with XAV-939 treatment (Be, Bk, Bq). CTR, control; COA, *Ctnnb1_+/flox(ex3)_ AMH-Cre*. See also Figure S6.

## Discussion

It has been demonstrated that the sexual fate commitment of germ cells is dependent on the differentiation of gonad somatic cells during sex determination. In female gonads, germ cells enter meiosis right after sex determination (McLaren, 1984), whereas the male germ cells are arrested in G0/G1stage and enter meiosis after birth (Western et al., 2008). Mesonephros derived RA is the most important extrinsic factor for meiosis initiation, germ cells won’t start meiosis in the absence of RA (Bowles et al., 2006; Koubova et al., 2006). It is widely believed that meiotic arrest of male germ cells during embryonic stage is mainly due to the expression of RA oxidizing enzyme CYP26B1 in Sertoli cells (Li et al., 2009). Interestingly, in this study, we found that disruption of testicular cords or depletion of Sertoli cells in testes did not lead to meiosis initiation of male germ cells during embryonic stages, suggesting that only RA is not sufficient to initiate meiosis. There must be some other factors cooperate with RA to regulate the meiosis initiation of germ cells.

The abnormal meiosis of male germ cells during embryonic stage is observed in several mouse models with male-to-female sex reversal. In *Sry* knockout males, germ cells are transformed into female germ cells with normal fertility (Kashimada and Koopman, 2010). In *Cyp26b1* and *Nanos2* knockout male mice, the transcription of *Stra8* and *Sycp3* is significantly increased in germ cells, yet the synaptonemal complex is not formed in *Nanos2* knockout mouse (MacLean et al., 2007; Suzuki and Saga, 2008), which is a typical marker of meiosis initiation. On the other hand, the defect of female germ cell meiosis was reported after inactivation of *Smad4* (Wu et al., 2016). A recent *in vitro* study also demonstrates that BMPs and RA act synergistically to specify female germ cell fate of *in vitro* differentiated PGC like cells under a defined condition (Miyauchi et al., 2017). All these studies suggest that the germ cell fate could be converted during sex determination. However, whether the fate of germ cells could be reprogrammed after sex determination remains an opening question.

In the present study, we found that the expression of meiosis associated genes was significantly increased in male germ cells by over-activation of CTNNB1 in Sertoli cells after sex determination. Moreover, synaptonemal complex was well organized and the germ cells at pachytene and diplotene stage were observed, indicating the male germ cells ectopically initiate meiosis after sex determination. Since Sertoli cells are transformed into granulosa-like cells in this mouse model, we also tested whether the male germ cells are transdifferentiated into female germ cell fate. The expression of several female germ cell marker gene was upregulated. However, the male germ cells were not clustered with control female germ cells by PCA analysis, indicating that the germ cells are not transdifferentiated into female cell fate. Previous study demonstrates that high levels of *Bmp2* and *Bmp5* in pre-granulosa cells is most likely instructive in specifying the female germ cell fate (Jameson et al., 2012b; Miyauchi et al., 2017). In this study, we found that both BMP2 and BMP4 are involved in inducing germ cell meiosis, and BMP4 shows more effects in this process.

*Dazl* is an important intrinsic factor which is essential for germ cell meiosis initiation, and germ cells will not start meiosis in the absence of *Dazl* (Gill et al., 2011). *Dazl* is expressed in both male and female PGCs and essential for female germ cell meiosis “license” (Lin et al., 2008). Previous studies find that *Nanos2* represses male germ cell meiosis by directly binding to 3’UTR of *Dazl* and causes the degradation of *Dazl* mRNA (Fukuda et al., 2018; Kato et al., 2016). In this study, we found that *Dazl* expression was significantly increased in the germ cells of COA mice and was reduced with BMP signaling inhibitor treatment. *In vitro* tissue culture experiments also confirmed that the expression of *Dazl* was induced by BMP signaling pathway, but RA couldn’t upregulate *Dazl* expression.

During germ cell development, the epigenetic modification undergoes global reconstruction (Hill et al., 2018) and previous study reported that *Dazl* is required for genome remethylating (Gill et al., 2011). However, the detailed regulation associated with germ cell meiosis is unclear. In this study, we found that DNA methylation was significantly decreased in the germ cells of COA testes and increased after BMP signaling inhibitor treatment. Moreover, the expression of DNA methyltransferase *Dnmt3a* was also decreased significantly in germ cells of COA testes and induced by BMP signaling pathway inhibitor. These results suggest that the DNA methylation of male germ cells during embryonic stage is most likely mediated by *Dnmt3a* and probably repressed by BMP signaling in female germ cells. However, we could not exclude other possibility which also regulated by BMP signaling pathway and need further investigation.

In summary, our study suggests that germ cell fate could be reprogrammed after sex determination when Sertoli cells were transformed into granulosa-like cells using *Ctnnb1* over-activated mouse. And BMP signaling pathway plays important roles in germ cell meiosis initiation. Moreover, our study demonstrates that BMPs are involved in inducing germ cell meiosis most likely by up-regulating *Dazl* expression and which is probably mediated by repressing DNA methylation. The results of this study provide more clues for better understanding the mechanism of germ cell meiosis initiation.

## Materials and Methods

### Mice

All animal work was performed according to the regulations of the Institutional Animal Care and Use committee of the Institute of Zoology, Chinese Academy of Sciences. All mice were maintained in a C57BL/6;129/SvEv mixed background. *Ctnnb1_+/flox(ex3)_ ROSA_mT/mG_ AMH-Cre* mice were obtained by crossing *Ctnnb1 _flox(ex3)/flox(ex3)_* (Harada et al., 1999)*; ROSA_mT/mG_* (Muzumdar et al., 2007) female mice with *AMH-Cre* transgenic males (Lecureuil et al., 2002). *DTA_+/flox_ AMH-Cre* mice were obtained by crossing *DTA_+/flox_* (Brockschnieder et al., 2004) females with *AMH-Cre* transgenic males. DNA isolated from adult tail tips and fetal tissues, and genotyping was performed by PCR as previously described (Gao et al., 2006; Harada et al., 1999).

For administering LDN-193189 into pregnant mice, LDN-193189 (S2618, Selleckchem) was dissolved in DMSO and then diluted in water and 2.5 mg/kg body weight was injected intra-peritoneally every 12 hr from E13.5 to E16.5.

### Tissue collection and histological analysis

The gonads were dissected at different developmental stages immediately after euthanasia, fixed in 4% paraformaldehyde for up to 24 hrs, stored in 70% ethanol at 4°C, and embedded with paraffin. Five-micrometer-thick sections were cut and mounted on glass slides. After deparaffinization, the sections were processed for Hematoxylin and Eosin (H&E) staining and immunofluorescent (IF) analysis. H&E stained sections were examined with a Nikon Microscope, and the images were captured with a Nikon DS-Ri1 CCD camera.

### Immunofluorescence analysis

After rehydration and antigen retrieval, 5 μm sections were incubated with 5% donkey serum in 0.3% triton X-100 for 1 hr. Then, the sections were immunolabeled with the primary antibodies for 1.5 hrs at room temperature. Antibodies were diluted as following: CTNNB1 (1:400, Abcam, ab6302), DDX4 (1:200, Abcam, ab5096), DAZL (1:200, Millipore, AB5535), STRA8 (1:200, Abcam, ab49405), SYCP3 (1:200, Abcam, ab15093), γH2AX (1:400, Millipore, 05-636) and GFP (1:100, Santa, sc-9996). After three times wash in PBS, the equivalent fluorescence labeled secondary antibodies (1:200, Jackson ImmunoResearch) were used for detection. DAPI was used to label the nuclei. After staining, the sections were analyzed with a confocal laser scanning microscope (Carl Zeiss Inc., Thornwood, NY).

### Chromosome spreads of germ cells

The meiotic prophase I stages were evaluated by chromosome spreading of germ cells from control ovaries and *Ctnnb1_+/flox(ex3)_ AMH-Cre* testes at E17.5. Briefly, tissues were disaggregated into single cell with 0.25% trypsin plus 0.2 g/L EDTA (Hyclone, SH3004202). After neutralization with 10% fetal bovine serum (FBS; Gibco, 10099-141), cells were treated in 1% trisodium citrate (Sigma, S1804) hypotonic solution for 20 mins at room temperature, then fixed in 1% paraformaldehyde (PFA, Beyotime, P0099). After spread onto poly-L-lysine (Sigma, P4707) precoated slides, and dried at 37°C, slides were blocked with ADB (3% BSA, 1% normal goat serum, and 0.005% Triton-100 in tris buffered saline, TBS) for 30 mins at room temperature and incubated overnight with 1: 400 diluted rabbit anti-SYCP3 antibody (1:200, Abcam, ab15093) at 37°C. After rinsing 3 times in TBS, incubated with 1: 400 diluted R-Phycoerythrin-conjugated goat anti-rabbit secondary antibody (Sigma, P9537) for 1.5 hrs at 37°C in dark. The meiotic prophase stages were determined under the Olympus BX51 fluorescence microscope with Cell Sens Ver.1.5 software by observing the characteristic patterns of SYCP3 immunostaining of the chromosomes.

### Nucleic acid isolation and quantitative reverse transcription-PCR

Total RNA was extracted using a Qiagen RNeasy kit (Qiagen, 74104) in accordance with the manufacturer’s instructions. Two micrograms of total RNA were used to synthesize first-strand cDNA. To quantify gene expression, a real-time SYBR Green assay was performed with isolated RNA. *Gapdh* was used as endogenous control. The relative level of candidate gene expression was calculated using the formula 2_-ΔΔCT_ as described in the SYBR Green user manual. The primers used for RT-PCR are listed in Table S1.

### Sertoli cell and single germ cell isolation

Control and CTNNB1 over-activated Sertoli cells from *ROSA_mT/mG_ AMH-Cre* and *Ctnnb1_+/flox(ex3)_ ROSA_mT/mG_ AMH-Cre* mice at E13.5 and E14.5 were isolated using flow cytometry based on GFP fluorescence as described previously (Li et al., 2017). The sorted cells were stored at −80°C for RNA extraction. Germ cells from control and *Ctnnb1_+/flox(ex3)_ AMH-Cre* mice at E14.5 and E15.5 were isolated as following. The dissected ovaries were washed with PBS for 3 times and digested with 0.05% trypsin-EDTA. After neutralization with 10% FBS, resuspended pellets to single cells and incubated with Anti-SSEA-1 (CD15) Microbeads (Miltenyi Biotec, 130-094-530). After collected the positive cells, single cell was picked out for RNA extraction by pipette under a dissecting microscope. After enzymatic dissociation, the solution was transferred into prepared lysis buffer with an 8-nt barcode.

### RNA-seq library construction and sequencing

Total RNA was extracted from isolated Sertoli cells of E13.5 and E14.5 embryos using Qiagen RNeasy kit in accordance to the manufacturer’s instructions. NEB Next Ultra RNA Library Prep Kit was used for RNA library construction. The RNA library was sequenced by Illumina Hiseq 2500 and aligned RNA-seq reads to Mus musculus UCSC mm9 references with the Tophat software (http://tophat.cbcb.umd.edu/), and the FPKM of each gene was calculated using Cufflinks (http://cufflinks.cbcb.umd.edu). The detailed procedure was performed as previously described (Yu et al., 2016).

### Single-cell cDNA amplification and library construction

Single-cell cDNA amplification was carried out using the STRT protocol as described previously (Dong et al., 2018; Fan et al., 2018). Libraries were prepared using KAPA Hyper Prep Kits (KK8505). The NEB U-shaped adapter was used for ligation. After 8–10 cycles amplification with primers, the libraries were sequenced with Illumina Hiseq2500 platform to generate 150-bp paired-end reads. To identify DEGs between control and CTNNB1 over-activated Sertoli cells, the number of unambiguous clean tags in each library was normalized to the TPM to obtain the normalized gene expression level. DEGs was identified as a false discovery rate (FDR) <0.001 and a threshold absolute log 2-fold change for the sequence counts across the libraries. Gene ontology and KEGG analysis was performed using DAVID Bioinformatics Resources 6.8.

### Gonads and re-aggregated tissues in vitro culture

The genital ridges were cultured on agarose blocks as described previously (Coscioni et al., 2001; Zhang et al., 2015b). The agarose block was made with 2% agarose and pre-balanced with culture medium up to 12 hrs. The fetal ovaries and testes from E11.5 and E13.5 embryos were dissected and placed on the pre-balanced agarose blocks. The gonads were cultured at 37°C in a saturated humidity incubator infused with 5% CO_2_ in air. For *in vitro* treatment, LDN-193189 (S2618, Selleckchem), XAV-939 (S1180, Selleckchem) and Kartogenin (S7658, Selleckchem) was dissolved in DMSO and used with a respective concentration of 10 μM.

The genital ridges from female and male embryos were separated from mesonephro and dissociated with 0.05% trypsin-EDTA at 37°C for 10 mins. After neutralization with 10% FBS, wash twice with DMEM supplemented with 10% FBS. Large clumps of cells were removed using a 70 μm cell strainer (Corning, 352340). Gonadal cells suspension was incubated with Anti-SSEA-1 (CD15) Microbeads (Miltenyi Biotec, 130-094-530). After washing, cell suspension with PBS supplemented with 0.5% BSA and 2 mM EDTA was applied to an MS column (Miltenyi Biotec, 130-042-201). After negative cells flow through the column, magnetically labeled cells were collected as the SSEA1-positive germ cells. The cells in the flow-through were used as gonadal somatic cells. Re-aggregated germ cells and somatic cells with the ratio of 1:10 and cultured on pre-balanced 2% agarose blocks at 37°C in a saturated humidity incubator infused with 5% CO_2_ in air.

### Statistical analysis

Experiments were repeated at least three times independently. Three to five genital ridges at each culture condition were used for immunostaining. The quantitative results are presented as the mean ± SD. The data were evaluated for significant differences using Student’s t-test and one-way ANOVA. P-values < 0.05 were considered to be significant.

## Acknowledgements

This work was supported by National key R&D program of China (2018YFA0107702); The National Science Fund for Distinguished Young Scholars (81525011); Strategic Priority Research Program of the Chinese Academy of Sciences (XDB19000000); and The National Natural Science Foundation of China (31601193, and 31671496).

## Supplemental information

**Figure S1.**
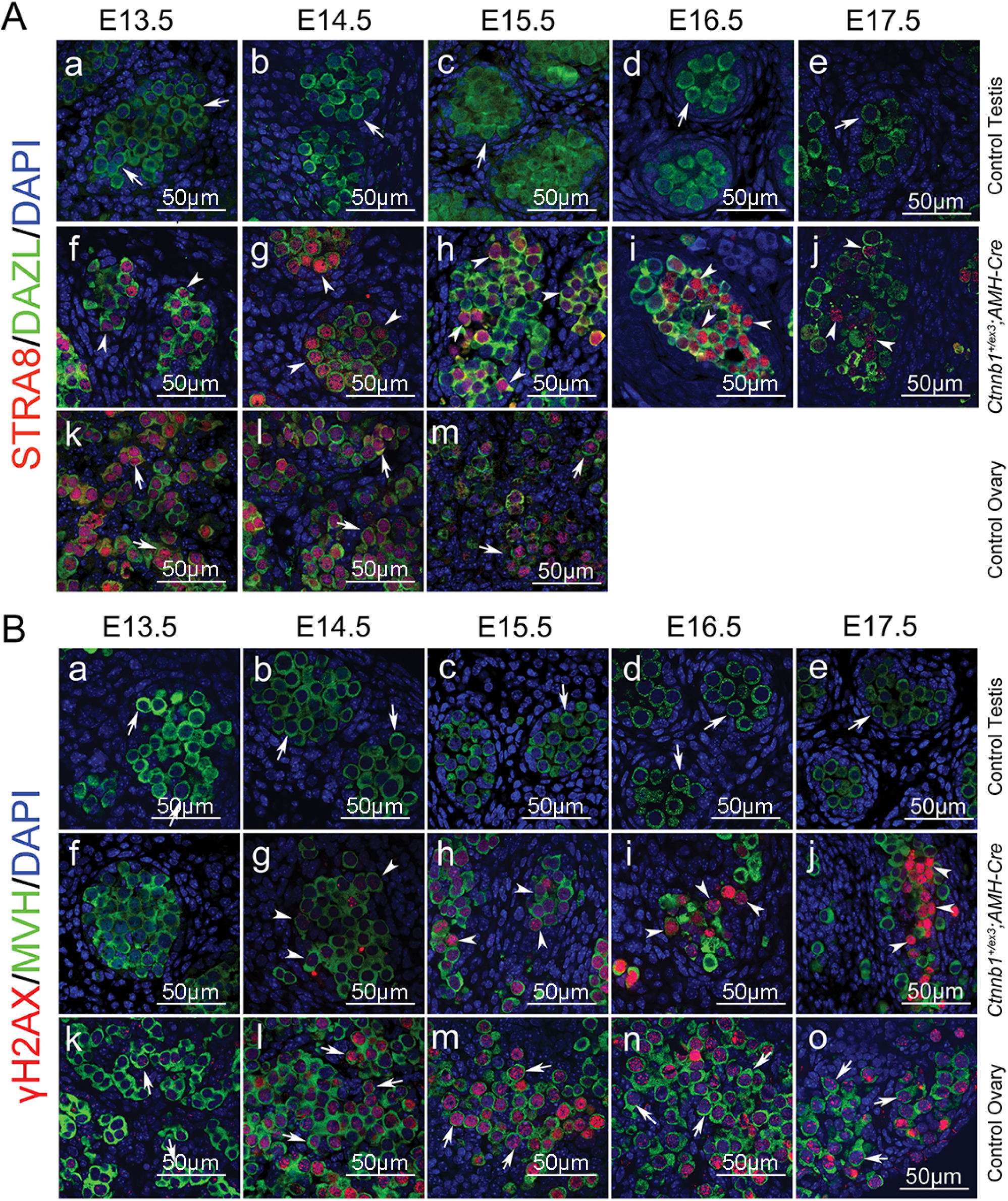
A. STRA8 and γH2AX were abnormally expressed in germ cells of *Ctnnb1_+/flox(ex3)_ AMH-Cre* testes during embryonic stage. **A.** No STRA8 signal was detected in the germ cells of control testes from E13.5 to E17.5 (Aa-Ae, white arrows). STRA8 was expressed in a large number of germ cells in *Ctnnb1_+/flox(ex3)_ AMH-Cre* testes from E13.5 to E16.5 (Af-Aj, white arrowheads). Most germ cells in control ovaries from E13.5 to E15.5 were STRA8-positive (Ak-Am, white arrows). **B.** No γH2AX signal was detected in MVH-positive germ cells of control testes from E13.5 to E17.5 (Ba-Be, white arrows). γH2AX-positive germ cells were noted in germ cells of *Ctnnb1_+/flox(ex3)_ AMH-Cre* testes from E14.5 to E17.5 (Bg-Bj, white arrowheads). Most of germ cells in control ovaries from E14.5 to E17.5 were γH2AX-positive (Bl-Bo, white arrows). Related to Figure 1.

**Figure S2.**
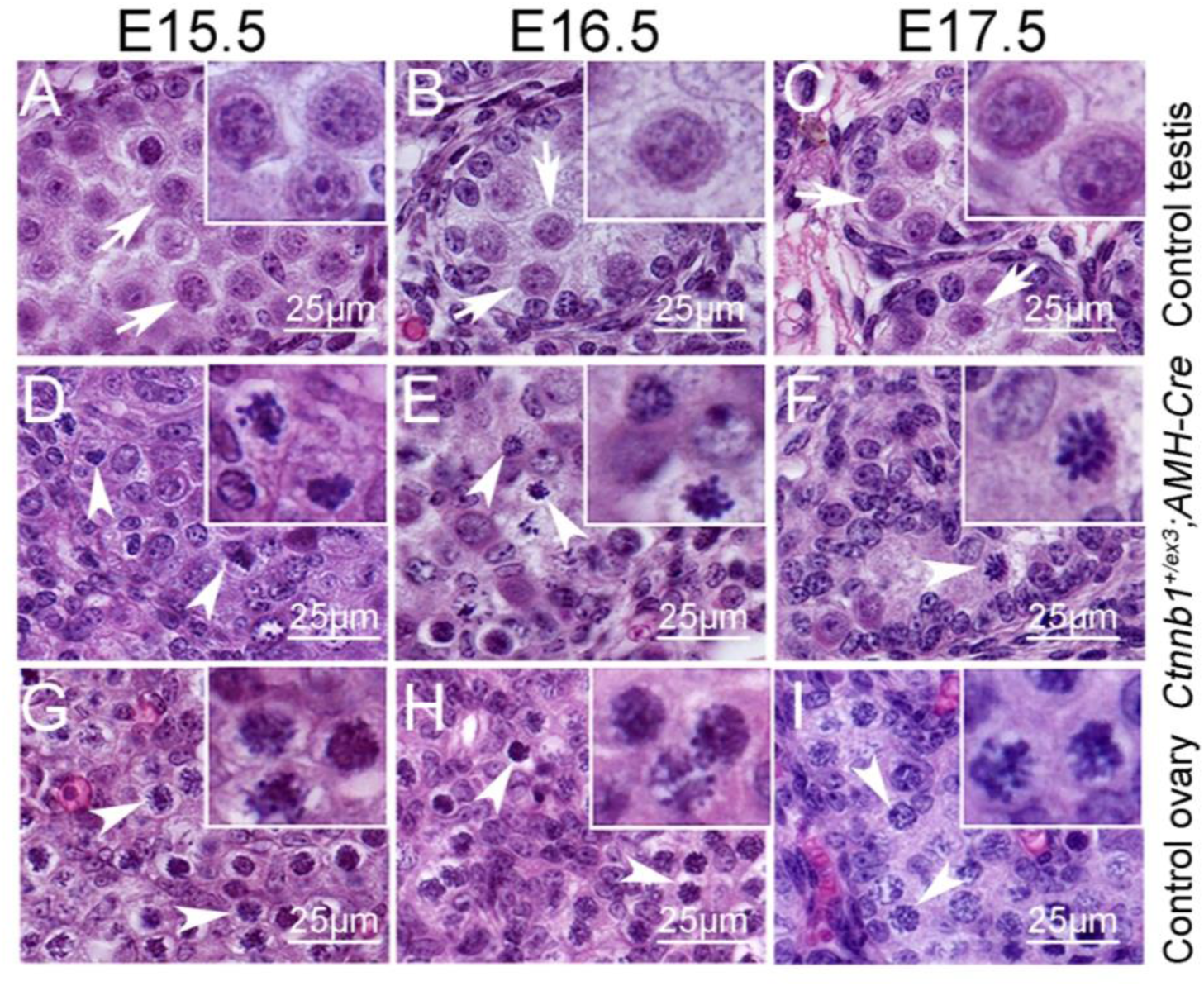
Chromatin condensation was observed in the germ cells of *Ctnnb1_+/flox(ex3)_ AMH-Cre* testes. No chromatin condensation was observed in germ cells of control testes from E15.5 to E17.5 (A-C, white arrows). The thread–like chromosome condensation was observed in germ cells of *Ctnnb1_+/flox(ex3)_ AMH-Cre* testes (A-F, white arrowheads), which was similar to the morphology of chromatin in germ cells of control ovaries (G-I, white arrowheads). Related to Figure 2.

**Figure S3.**
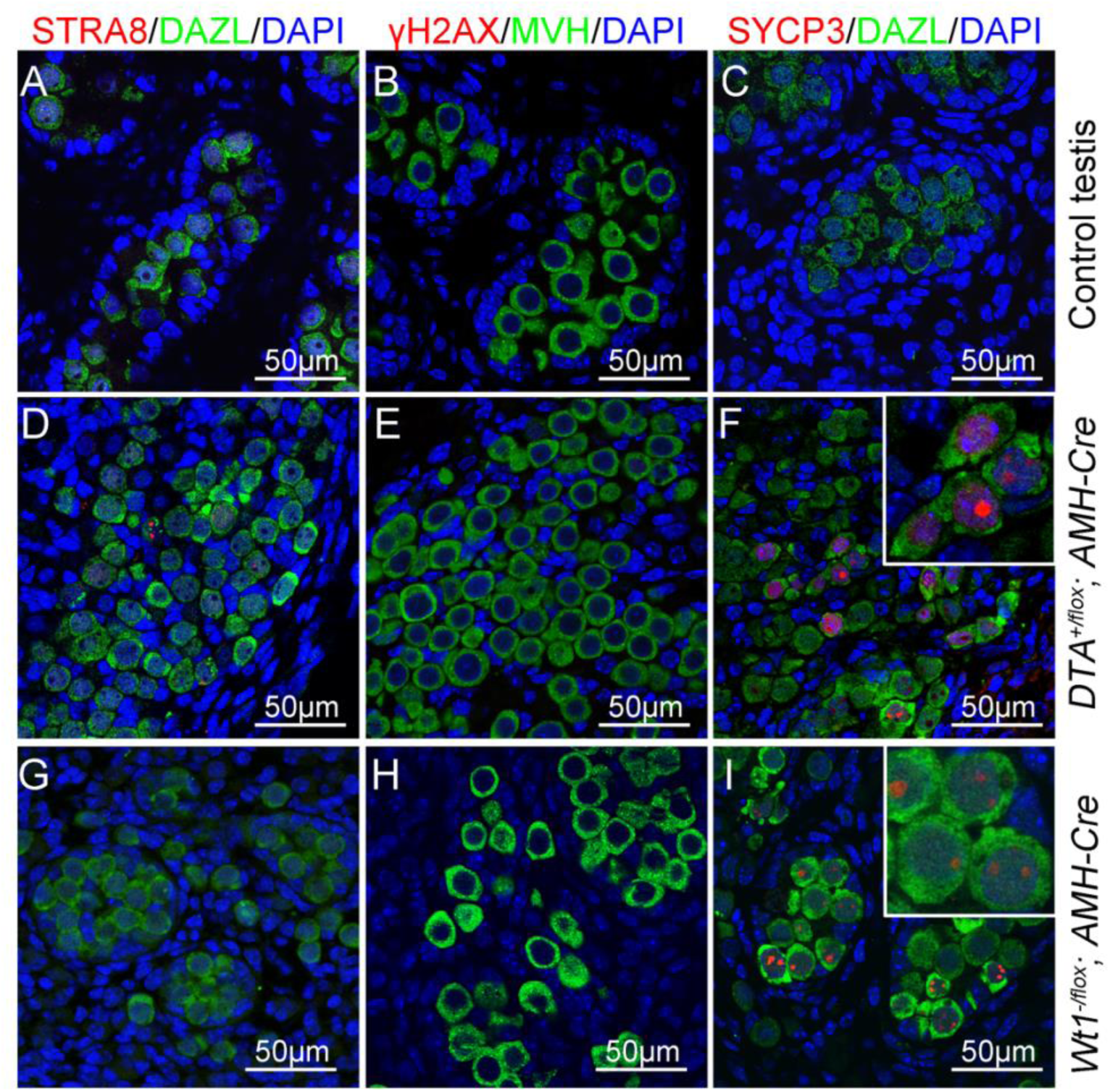
No meiotic germ cell was observed in testes of *DTA_+/flox_ AMH-Cre* and *Wt1_-/flox_ AMH-Cre* testes at E16.5. Germ cells in control testes were negative for STRA8 (A), γH2AX (B) and SYCP3 (C) proteins. No STRA8 (D and G) and γH2AX (E and H) signal were detected in germ cells of *DTA_+/flox_ AMH-Cre* and *Wt _-/flox_ AMH-Cre* testes. Weak SYCP3 signal was detected in germ cells from *DTA_+/flox_ AMH-Cre* (F, inset) and *Wt _-/flox_ AMH-Cre* testes (I, inset).

**Figure S4.**
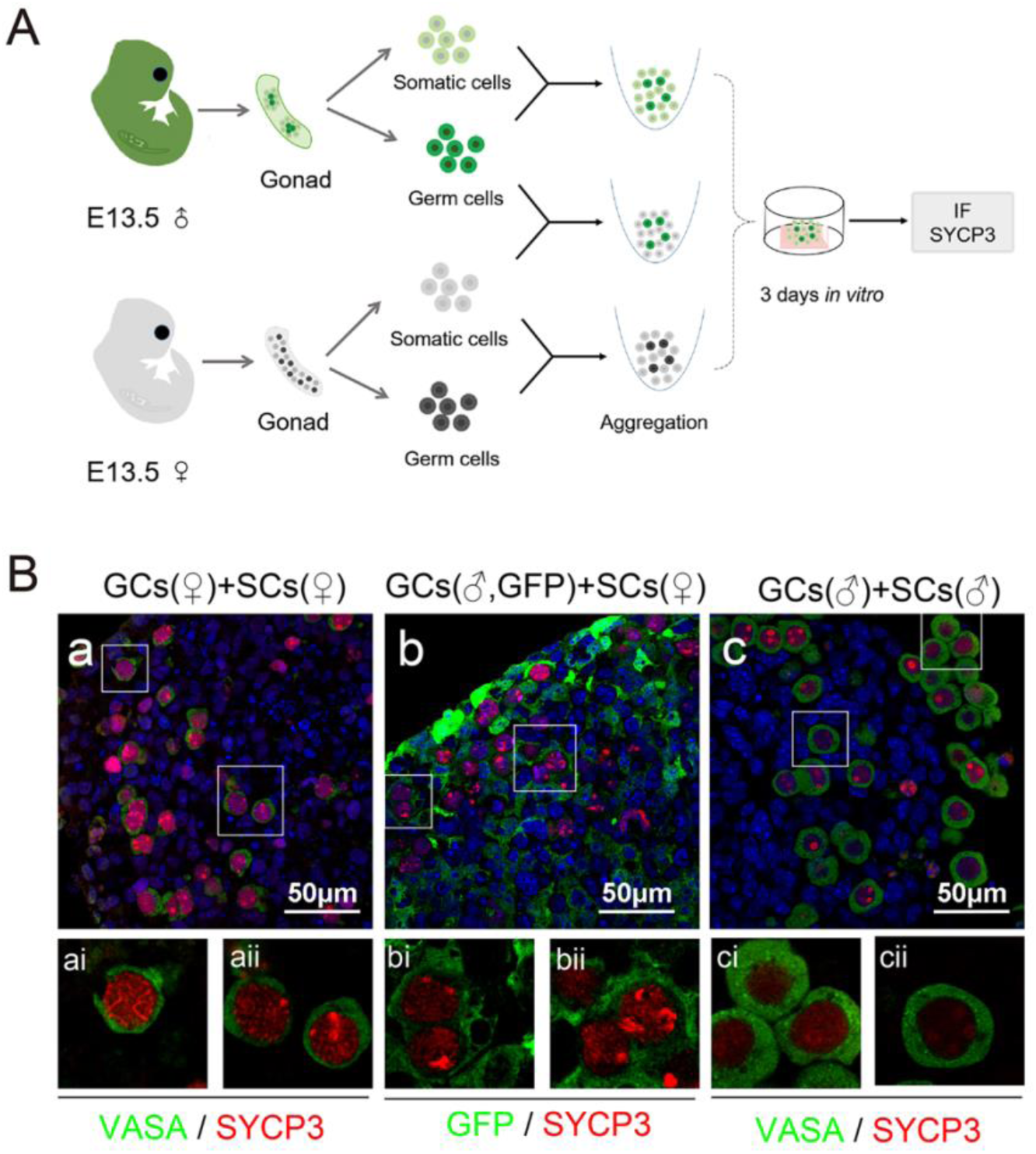
Meiosis in embryonic male germ cells was induced by female somatic cells. **A.** The schematic diagram of re-aggregation and culture of germ cells (GCs) and somatic cells (SCs) from E13.5. **B.** Thread-like SYCP3 signal was observed in female germ cells aggregated with female somatic cells (Ba). Thread-like SYCP3 was noted in the male germ cells (GFP_+_, green) aggregated with female somatic cells (Bb). Very weak SYCP3 signal was detected in the aggregation of male germ cells with male somatic cells (Bc).

**Figure S5.**
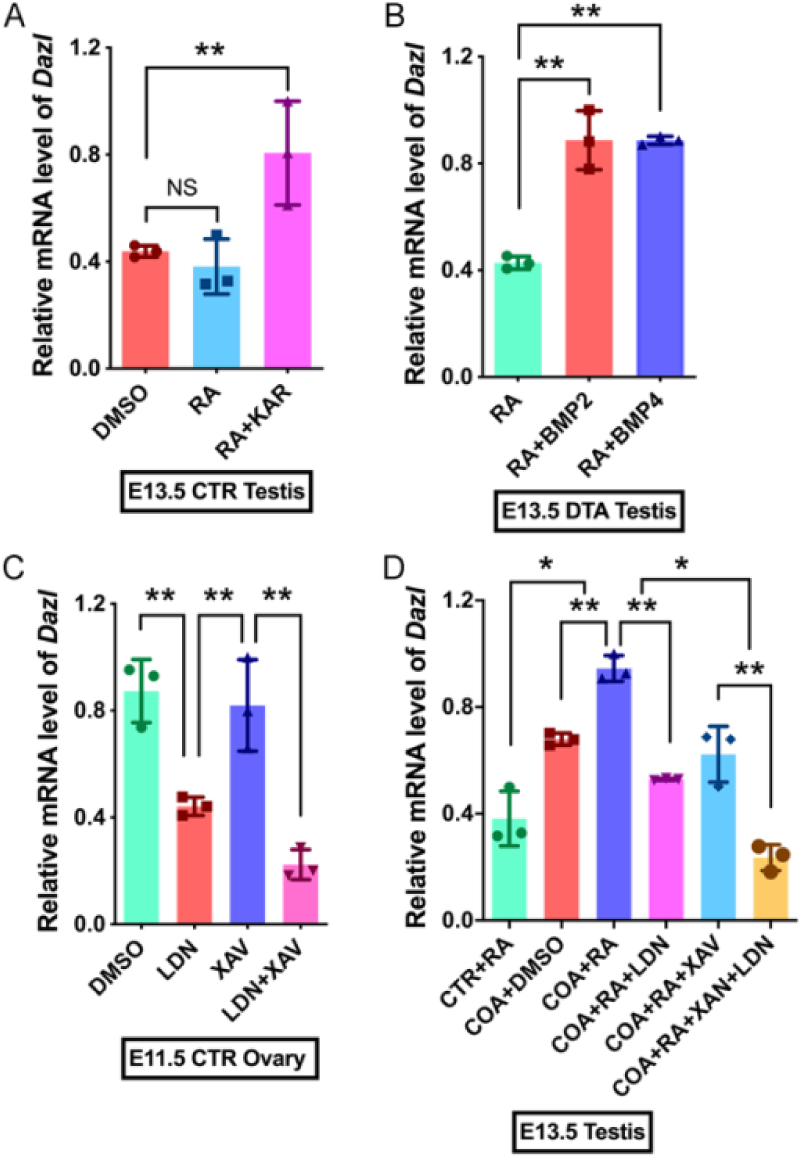
The mRNA level of *Dazl* was induced by BMP signaling. **A.** The expression of *Dazl* was significantly increased in KAR treated testis. **B.** The expression of *Dazl* was significantly increased in cultured *DTA_+/flox_ AMH-Cre* testis with BMP2 and BMP4 treatment. **C.** The expression of *Dazl* in control ovary was repressed by LDN-193189. **D.** The expression of *Dazl* in COA testis was inhibited by LDN-193189 and XAV-939 treatment. Data are presented as the mean ± SD. The student‘s unpaired t-test and Two-way ANOVA was used to test significance (*P< 0.05; **P< 0.01; ns: no significance). CTR, control; COA, *Ctnnb1_+/flox(ex3)_ AMH-Cre*.

**Figure S6.**
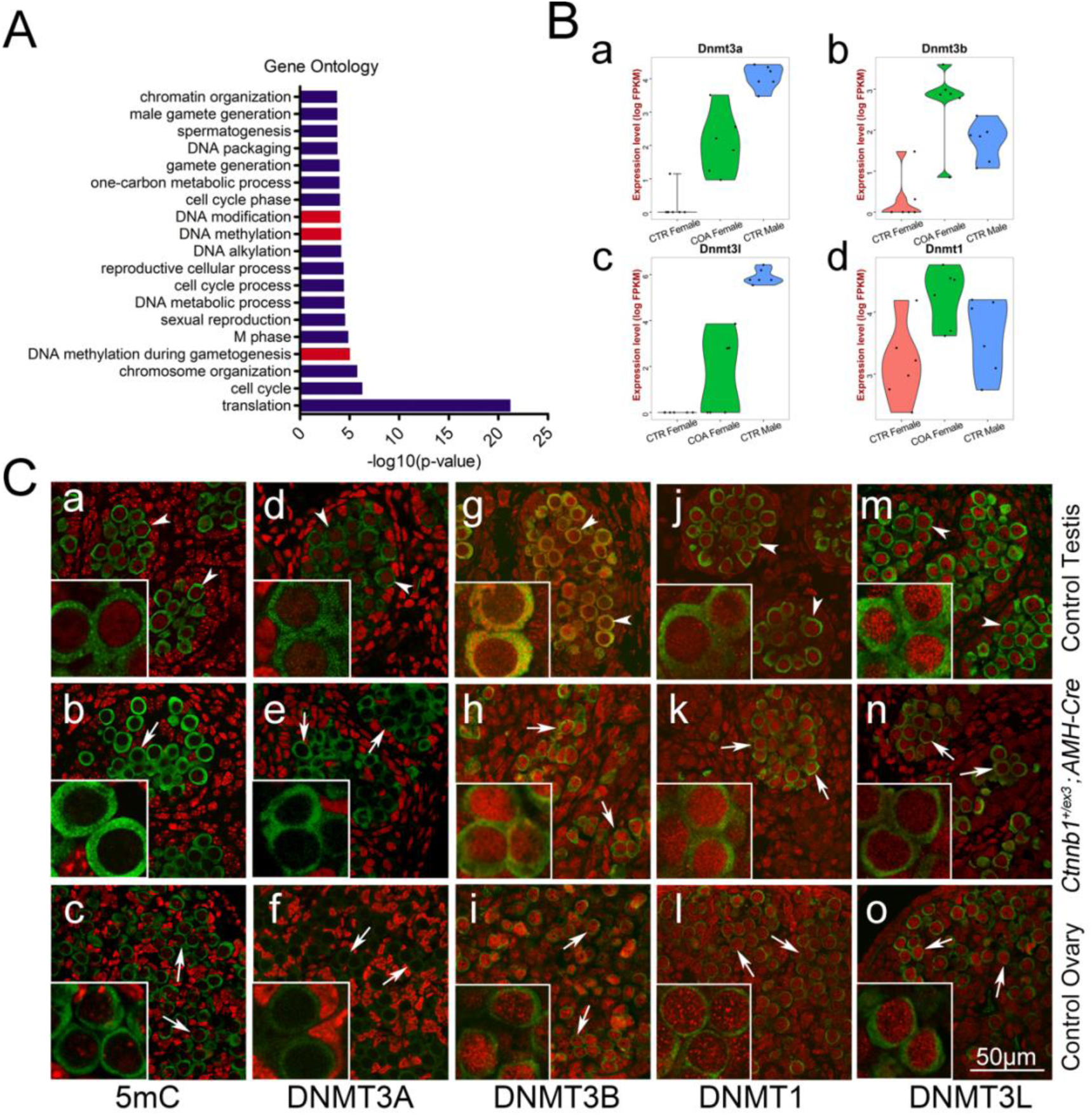
The status of DNA methylation was changed in germ cells of *Ctnnb1_+/flox(ex3)_ AMH-Cre* mice. **A.** Gene ontology analysis of DEGs at E15.5. **B.** Expression level of DNA methyltransferase (*Dnmt3a*, *Dnmt3b*, *Dnmt3l*, *Dnmt1*) at E15.5 in control male, COA male and control female germ cells. **C.** Immunostaining of 5mC and DNA methyltransferase. CTR, control; COA, *Ctnnb1_+/flox(ex3)_ AMH-Cre*. Related to Figure 6.

**Table. S1.**
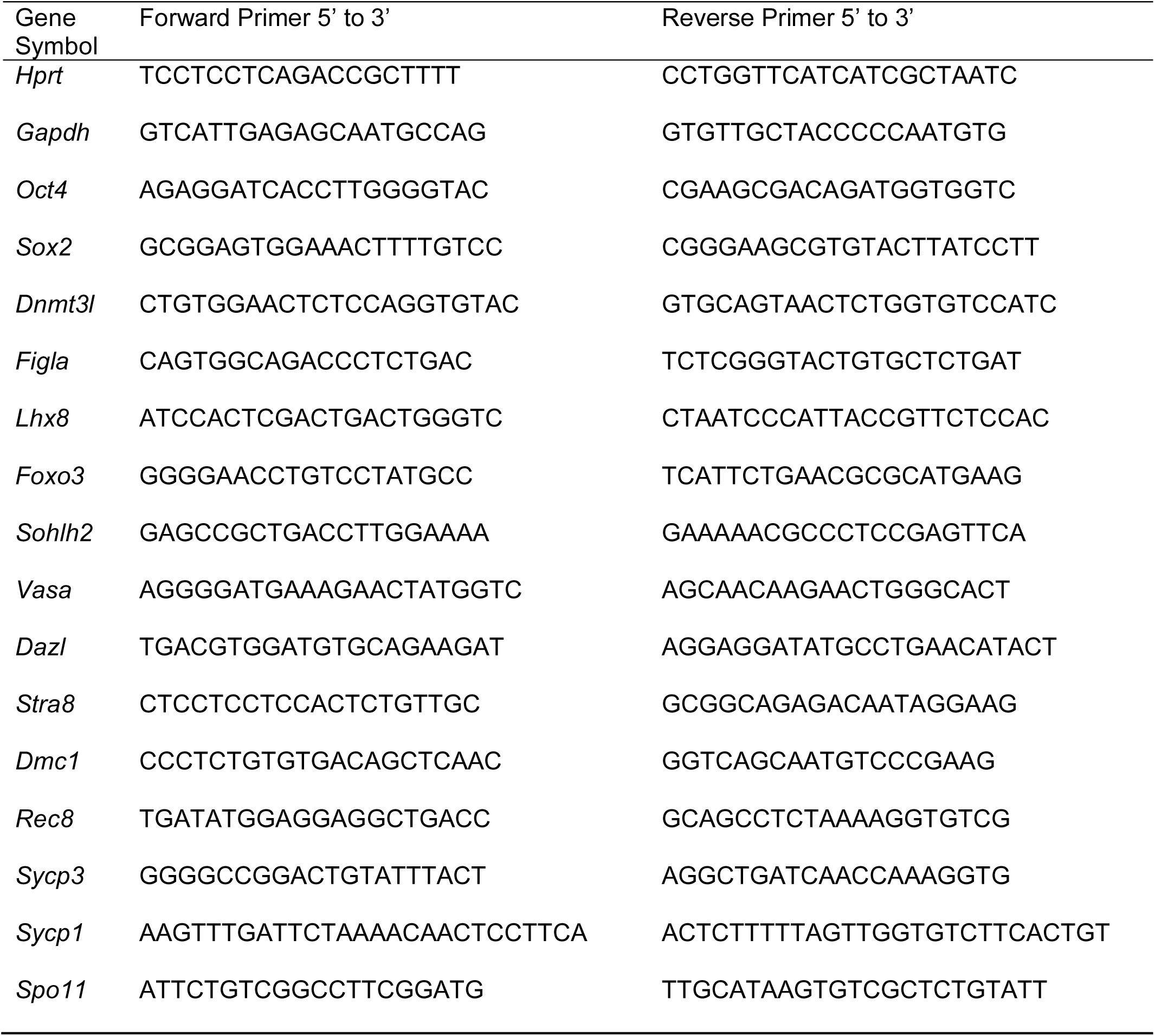
Primers for quantitave RT-PCR

Table. S2. Somatic cells sequencing data and analysis related to Figure 2.

Table. S3. Single germ cell sequencing data and analysis related to Figure 3.

